# Aggressive conflict as a transient and distributed landmark in homing desert ants

**DOI:** 10.1101/2024.11.27.625616

**Authors:** Antonio Bollig, Marília Freire, Konrad Bücking, Jana Kühnapfel, Markus Knaden

## Abstract

Across many species of the animal kingdom, the ability to navigate long distances is important for securing food and other resources. For example, ants, bees, and rats have long been known to use path integration, a mechanism to integrate distance and direction travelled, to find their way back to their nest. However, path integration is imperfect, and errors accumulate over time as animals forage. There are several ways to counteract this uncertainty, for example by utilizing local olfactory or visual landmarks along a familiar route to infer an individual’s position relative to its nest, i.e. its homing vector. Local landmarks are thought to be effective if they are constant, granting a strong association with the discrete location it is located at. Here we show that ants can utilize something much sparser and more distributed than an individual constant landmark to inform its homing vector. Our data suggests that Tunisian desert ants integrate intraspecific contact, or social cues, to augment navigation. We demonstrate that contact with foreign conspecifics at a position where the nest is expected diminishes path integration certainty, thereby expanding the nest search pattern. Furthermore, we show that ants form associations between intraspecific conflict and discrete locations to create so called local vectors, within which the homing path to the nest can be stored.

**Highlights:**

- Contact with foreign ants reduces path integrator certainty and alters homing strategy
- Pattern of systematic search could be altered by persistent adaptions of walking speed
- Foraging ants can form local vector memories using intraspecific conflict as a landmark

## Introduction

The desert ant *Cataglyphis fortis* inhabits the open salt pans of Tunisia, where it individually forages for dead arthropods killed by the heat. During their extensive foraging runs that can cover more than one kilometre and last for more than an hour (Buehlmann et al., 2014; Freire et al., 2023; Huber & Knaden, 2015) the ants use path integration (PI) to finally return to their mostly inconspicuous nest entrance. By combining a step integrator (Wittlinger et al. 2006) and a skylight compass (Wehner & Muller, 2006) the ants continuously compute their position relative to the nest entrance, and upon finding a food item, follow a PI vector that leads them to the starting point of their search, i.e. the nest entrance. Therefore, a reeled-off PI vector usually means that the nest is close. However, PI is error prone, and errors accumulate with increasing walking distances (Müller and Wehner 1994). As a result, it does not necessarily guide the ant to the nest entrance directly but rather to its close vicinity. If a returning forager does not immediately find its nest entrance at this point, it will engage in systematic search behaviour (Mueller & Wehner, 1994). During this search, the ant runs loops of increasing size while repeatedly returning to the location where the PI indicated the nest position. The area covered by this search depends on the ant’s PI certainty, i.e. the ant’s trust in the accuracy of its path integrator, with ants returning from longer distances performing a wider search to compensate for accumulated errors (Merkle et al., 2006). Colonies of *C. fortis* are generally hostile toward one another yet are often located less than 10 m apart. Avoiding foreign colonies while exploiting possible social cues during homing poses a serious challenge to the foragers. Our study provides insight into how compensatory behaviours, such as systematic nest search, can be augmented and adapted through simple behavioural computations, and how minor adjustments can modify complex navigational algorithms essential for the survival of these resourceful extremophiles.

In the first part of this study, we use the width of the systematic search to quantify the ants’ certainty in their PI vector and investigate whether this certainty is affected by social cues. Specifically, we test whether contact with nestmates just before the beginning of the systematic nest search leads ants to assume close nest proximity and thus exhibit a narrower systematic search, and whether contact with foreign ants has the opposite effect, reducing certainty and widening the search pattern. In the second part, we explore whether strongly distributed landmarks like places of frequent intraspecific conflict can be associatively tied to local vectors indicating the relative position of the forager to its nest entrance. We test whether extended exposure of foragers to an active conflict zone results in a retrievable vector memory indicating the foragers’ nest entrance.

## Results and discussion

We hypothesized that contact with nestmates at the beginning of the search would increase the ants’ certainty in the PI-defined nest position and therefore result in a narrower search, whereas contact with ants from a foreign colony would decrease this certainty and result in a wider search. After exposure to nestmates, ants exhibited a search pattern that did not differ significantly from that of control ants (Fig. 1c, blue vs grey). This suggests that contact with nestmates upon arriving in the vicinity of the nest entrance is not sufficient to increase the ants’ certainty in the PI-defined position of the nest entrance under the conditions tested. In contrast, interactions with foreign ants altered the nest search behaviour: enemy-exposed ants showed a significantly wider search pattern than control ants (Fig. 1c, red vs grey), suggesting a reduction in certainty about the PI-defined nest position following encounters with foreign ants.

**Fig. 1.**
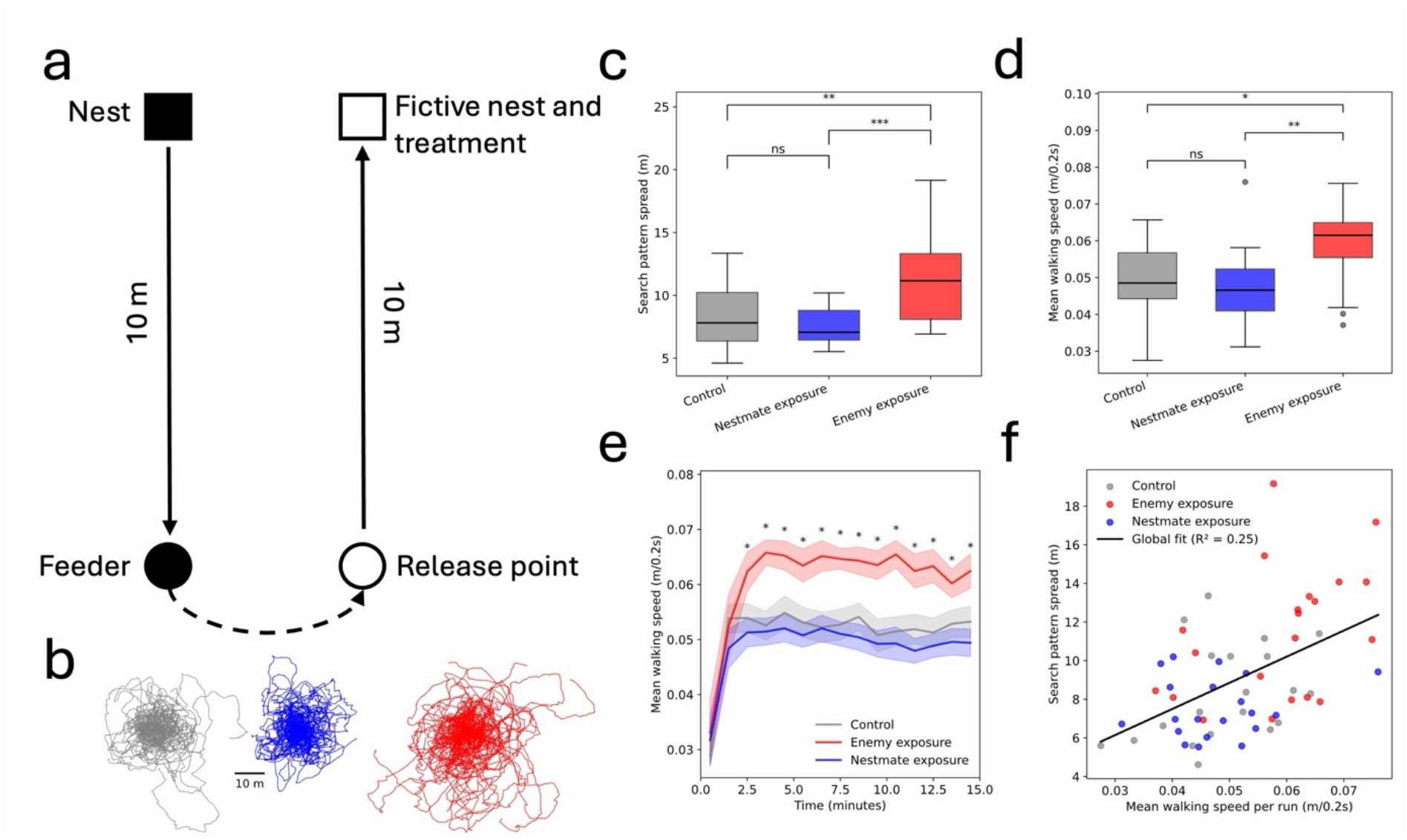
Contact with foreign conspecifics reduces PI certainty. (a) Experimental paradigm. Ants were displaced from a feeder (dashed arrow) to a release point and allowed to reel off their home vector toward the fictive nest position. Upon reaching this position, ants were exposed to one of three treatments: no social contact (control; colour-coded in grey), contact with nestmates (colour-coded in blue), or contact with foreign ants (colour-coded in red). (b) Representative overlaid systematic search trajectories for all ants in each treatment, colour-coded per treatment. (c) Search pattern spread across treatments, quantified as the radius of gyration of search trajectories, colour-coded per treatment (Kruskal–Wallis test followed by Mann–Whitney U tests). (d) Mean walking speed per ant run across treatments, colour-coded per treatment (Kruskal–Wallis test followed by pairwise Mann–Whitney U tests). (e) Development of walking speed during the systematic search. Lines show mean ± SEM; asterisks indicate time bins with significant differences between treatments (Kruskal-Wallis test). (f) Relationship between walking speed and search pattern spread across all runs; points are colour-coded by treatment (linear regression and Pearson correlation). Statistics legend: ns = not significant; *p < 0.05, **p < 0.01, ***p < 0.001).

Further examination of the ants’ runs, specifically locomotion speed, provided insight into how such behavioural changes may arise. Ants exposed to foreign ants showed a persistent increase in walking speed compared to both control ants and ants exposed to nestmates (Fig. 1d). This significant difference in walking speed was sustained throughout the 15-min search period and was not limited to an initial phase following exposure (Fig.1e). Across all runs, walking speed was positively correlated with search pattern spread (Fig. 1f). Together, data suggests that adjustments in locomotion speed may contribute to the modulation of systematic search behaviour in response to social stimuli.

The shape and spatial extent of systematic search behaviour are known to depend on search duration (Merkle et al., 2006). Whether the size and structure of a systematic search pattern can also be modulated by walking speed has, to our knowledge, not been directly tested. From an ecological perspective, it is plausible that the pattern of a systematic search might be tied to distance travelled rather than time passed, as foragers can differ substantially in size and locomotor speed. Future experiments, for example by shortening or elongating ants’ legs, will be required to determine whether distance travelled rather than time elapsed determines the structure of systematic search patterns.

In resource dense environments ant colonies can co-occur within less than 10 meters. With ants of other colonies being in proximity and fights between colonies being frequent (Buehlmann et al., 2012; Knaden & Wehner, 2004), distributed landmarks such as hostile encounters could be used to facilitate homing. Hence, we asked whether ants could retain spatial information about locations where such aggressive encounters occurred, potentially allowing them to infer their relative nest direction and distance (i.e. establishing a local vector pointing from the area of conflict to the nest). To test this, ants from two neighbouring nests were trained to a shared feeding zone, where repeated aggressive interactions with foreign ants occurred (Fig 2a).

**Fig. 2.**
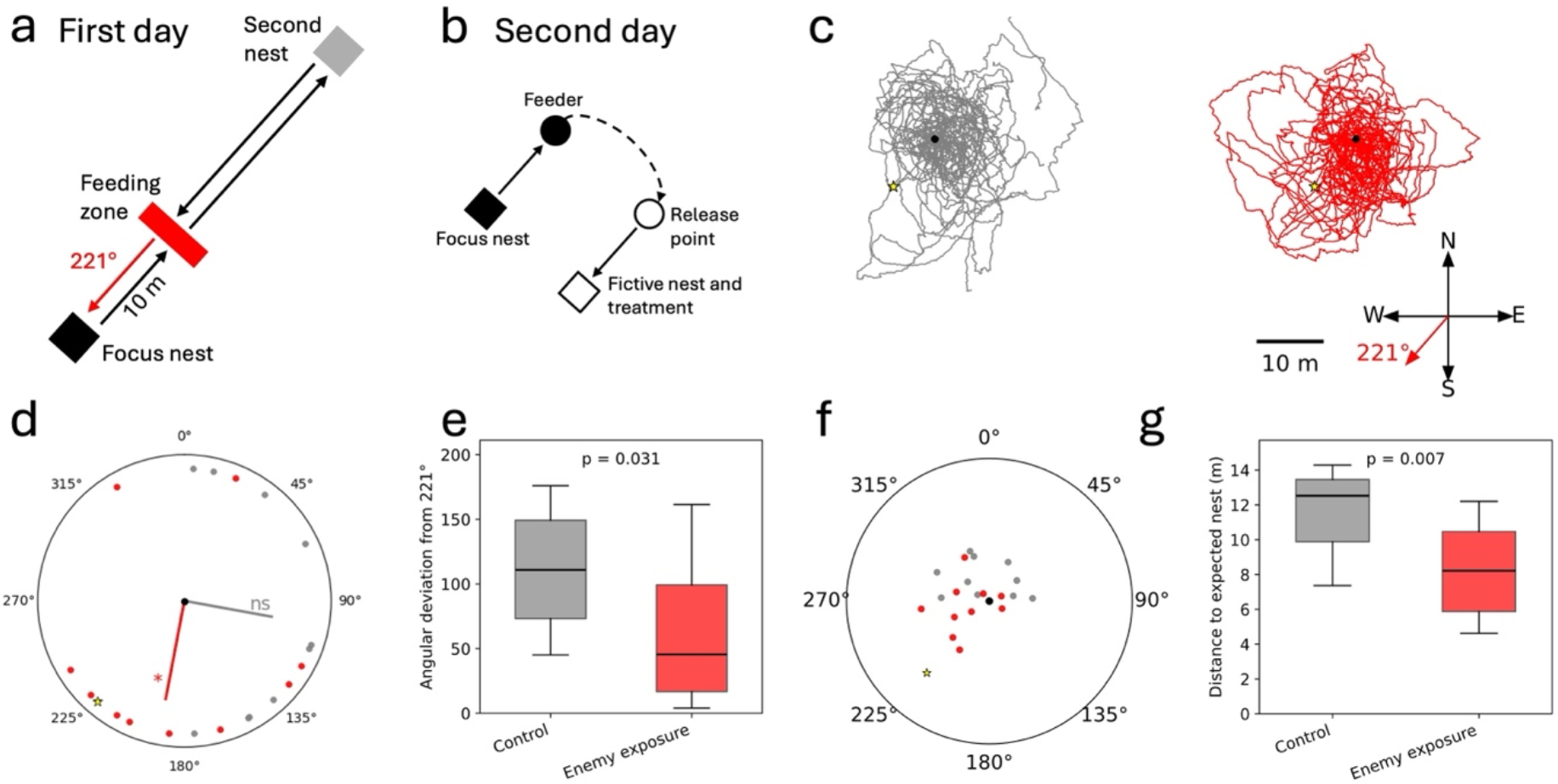
Desert ants form local vectors using areas of intraspecific conflict as a landmark. (a) Experimental paradigm on the first day: ants from two nests were trained to a shared feeding zone. (b) Experimental paradigm on the second day: ants from focal nest were trained to a single feeder in the same direction as the former shared feeding zone. (c) Overlaid search patterns of all ants across treatments. Black dots indicate starting positions; yellow stars mark the expected position of the nest based on training. (d) Circular plots showing centroid directions of systematic searches relative to the expected nest direction as indicated by yellow star. Lines indicate mean directions for each treatment, with significance of non-uniformity and clustering around expected direction indicated. (one-sample circular V-tests) (e) Angular deviation of centroid direction from the expected nest direction across treatments (Two-Sample Permutation test). (f) Spatial distribution of search centroids. Black dot indicates starting position of runs; yellow star marks the expected nest position according to training. (g) Euclidean distance of search centroids to the expected nest position across treatments (Two-Sample Permutation test). Statistics legend: ns = not significant; *p < 0.05, **p < 0.01, ***p < 0.001).

It is known that ants can recognize ants from colonies with which they have had aggressive interactions (Knaden & Wehner, 2003). Thus, we hypothesized that the focal ants might form experience-dependent spatial associations linking this zone of intraspecific conflict with a local vector pointing to their relative nest position. If so, exposure to foreign ants should bias their systematic search toward the nest direction associated with the former shared feeding zone (221° SW, Fig. 2a). We, therefore, let ants that just returned from the conflict area run off their home vector, exposed them to enemy ants from the conflict area, and let them begin their systematic search in a remote test field. Consistent with our hypothesis, the enemy-exposed ants showed a significant shift in search centroid toward the predicted nest direction compared to non-exposed control ants (Fig. 2d,e). In addition, the centroids of the enemy-exposed ants were, on average, located closer to the expected nest position defined by training (Fig. 2f,g).

Together, these results indicate that foraging ants of *C. fortis* use intraspecific cues to adjust their nest search strategy, but that such adjustments occur specifically in an aggressive context. Exposure to nestmates alone did not significantly narrow the search pattern and, hence, it is not sufficient to increase forager’s path integrator certainty (Fig. 1c). In contrast, encounters with foreign ants at the assumed nest entrance reduced PI certainty and led to broader nest searches (Fig. 1c). At the same time, ants that had previously experienced repeated conflict with a neighbouring colony appeared capable of forming spatial associations related to that conflict and recalled this information upon renewed contact with foreign ants, biasing their systematic nest search toward the direction associated with the former area of conflict (Fig. 2a). This behaviour is consistent with the idea that ants may use experience-dependent intraspecific cues as a form of local landmark during homing.

Path integration has previously been shown to influence aggression in *C. fortis*, with ants that perceive themselves as close to home displaying higher aggression levels than ants that perceive themselves far from home (Knaden & Wehner, 2004). Our results suggest that the relationship between aggression and path integration is bidirectional: aggressive social interactions can also influence PI-related behaviours. Encountering foreign ants at the beginning of the nest search reduced PI certainty and widened the search pattern, while prior experience with aggressive interactions allowed ants to bias their search based on “memorised” locations associated with conflict. This behaviour may be mediated by a nest-directed vector memory associated with the memorised location of conflict, similar to local vectors that *Catagylyphis* ants are known to associate with visual landmarks along familiar routes (Bisch-Knaden & Wehner, 2003; Collett et al. 1998). Incorporating local vectors facilitated by intraspecific cues as landmarks could help ants infer a corrected home vector on long homing bouts with erroneous path integration.

The existence of local vectors and global vectors in spatial navigation, as well as the tendency of local vectors associated with local landmarks inhibiting global vector expression on familiar routes, is well described in ants (Collet et al., 1998). Furthermore, the capacity of hymenopterans to integrate encounter rates into decision making is known to play a role in quorum sensing within the colony (Pratt 2005; Smith et al., 2017). Our study links these concepts by proposing that, in ants, local vectors can not only be tied to visual landmarks, but also to less permanent and more stochastic cues such as encounters with foreign ants.

*C. fortis* has been shown to judge the reliability of landmarks, i.e. to ignore visual or olfactory cues that are omnipresent and hence not place specific (Huber & Knaden 2017). Intraspecific social cues are inherently dynamic and unreliable compared to constant landmarks like rocks or brushes, suggesting that ants must flexibly weigh their influence against other navigational inputs such as path integration and landmark memory. This raises the possibility that the navigational system of *C. fortis* operates as a hierarchical or context-sensitive integration process, in which the reliability and ecological relevance of different cues determine their behavioural impact.

Future work should address the persistence and specificity of these “socially” derived spatial memories, as well as the mechanisms by which they are encoded and retrieved. In particular, determining whether such “social landmarks” generalize across contexts or remain tightly linked to specific locations and colony identities will be critical for understanding their role in navigation. More broadly, our results highlight the importance of considering sparse cues like social information as a possible component of spatial cognition, opening new avenues for studying how animals integrate cues to guide navigational behaviour.

## Methods

### Materials availability

This study did not generate any new unique reagents.

### Experimental Model and Study Participant Details

The model organism of this study is the desert ant *Cataglyphis fortis*. The desert ants inhabit a saltpan in Tunisia close to the village Menzel Chaker (Sebkhet Bou Jemel, 34°96’N, 10°41’E) and experiments were performed in May and June of 2023 and June of 2024. Only female foragers were used as subjects in this study.

### Experimental Design

In the first experiment, we established an artificial feeder (a shallow depression on the ground filled with biscuit crumbs) at 10 m from the nest (Fig. 1a). After approximately 20 min, the feeder was well established, with many ants travelling between nest and feeder. Individual ants were captured at the feeder using a falcon tube and marked with a spot of enamel paint on their gaster. After the paint had dried, ants were released at a remote test field approximately 100 m away from the feeder. Upon release, ants were provided with a food crumb and immediately reeled off their 10 m home vector (Fig. 1a). Ants that had reeled off their vector without deviating more than 1.5 m laterally were recaptured using a circular arena (diameter, 30 cm; height, 15 cm) placed around the ant. Within this arena, ants were assigned to one of three treatment groups. In Group 1 (control), the ant remained alone in the arena for 1 min. The arena was then lifted, allowing the ant to initiate its systematic search for the nest entrance. The search was recorded for the next 15 min using a GNSS antenna (GNSS-Commander; grey tracks in Fig. 1b,c). In the remaining treatments, additional ants were introduced to the arena to allow social interactions. Upon introduction, the test ant usually contacted these ants within a few seconds, resulting in intense antennation when interacting with nestmates and threatening behaviour or even escalating fights with foreign ants. In Group 2 (nestmate-exposed), the test ant was exposed in the arena to four nest mates, previously captured at their nest entrance, until it had made at least 10 direct antennal contacts with nest mates (on average, this took around one minute). Afterwards, the nest mates were removed, the test ant was released, and its systematic search recorded as described above (blue track in Fig. 1b). In Group 3 (foreigner-exposed), the procedure was identical to that of Group 2, except that the test ant was exposed to four foreign ants captured at the entrance of their foreign nest.

This usually resulted in aggression, including mandible threats and biting. After at least 10 such contacts, the foreign ants were removed, the test ant was released, and its systematic search was recorded as described above (red track in Fig. 1b).

In a second experiment, we examined whether ants retain spatial information associated with previous aggressive encounters. We selected a focal nest with a single neighbouring nest located approximately 100 m away (at a direction of 221°SW) and established a shared feeding zone (1.5 m x 3 m) between the two nests, positioned 10 m from the focal nest (Fig. 2a). Placing the feeding zone closer to the focal nest allowed us to regulate the influx of foreign ants while maximizing the number of foragers from the focal nest reaching the site. Biscuit crumbs were provided ad libitum at the feeding zone, resulting in exploitation by both colonies and frequent aggressive interactions (mostly mandible threats) between ants from the two nests. After one day of training, we removed the shared feeding zone and limited forager influx from the neighbouring nest. After a feeding break of three hours, we re-established a singular feeder at 10 m from the focal nest, in the same direction as the former feeding zone, but much smaller. This feeder was only exploited by ants of the focal nest. Then, we captured foraging ants at the feeder, displaced them to a remote test field, and allowed them to reel-off their PI vector. After, we captured ants using the circular arena, where they were either kept isolated for approximately 1 min (control) or exposed for the same time to four foreign ants from the colony they previously encountered at the shared feeding zone. Afterwards, the focal ants were released and their search for the nest entrance was recorded for 15 min.

## Quantification And Statistical Analysis

### Search Pattern Spread per Ant Run

Search pattern spread was quantified for each ant run based on the spatial distribution of its systematic search. For each NMEA track, implausible step jumps greater than 5 m (due to recording inaccuracies) were removed, and the radius of gyration (root-mean-square distance from the centroid) of all remaining GPS positions was calculated. Distribution of per-run spread across treatments are shown as boxplots, and overall and pairwise differences between groups were assessed using a Kruskal–Wallis test followed by Bonferroni-corrected Mann– Whitney U tests to account for multiple pairwise comparisons.

### Mean Walking Speed per Ant Run

Mean walking speed was quantified for each ant run based on the movement between consecutive GPS positions. Recording frequency of the GPS device was 5 Hz. Using the same processing steps as described above, step jumps greater than 5 m were removed from each NMEA track. GPS coordinates were converted to decimal degrees, and step distances between consecutive points were computed and averaged per run as a proxy for walking speed. Differences between treatments were assessed using a Kruskal–Wallis test with followed by pairwise Mann–Whitney U tests.

### Step Distance Development Over Time

Step distance development over time was quantified for each ant run based on movement between consecutive GPS positions. Using the same processing steps as described above, step jumps greater than 5 m were removed from each NMEA track. Step distances were calculated between consecutive positions, grouped into 60 second time bins over the 15 minutes search period, and averaged across runs to obtain mean ± SEM values per walking speed per treatment over time. Differences between treatments were assessed separately for each time bin using Kruskal-Wallis tests.

### Global Spread vs Walking Speed Correlation

The relationship between search pattern spread and walking speed was quantified across ant runs. For each run, mean walking speed (calculated as mean step distance after discarding steps jumps greated than 5 m) and search pattern spread (radius of gyration of cleaned GPS positions) were extracted as a pair of values. These values were pooled across treatments and analsyed using a linear regression and Pearson correlation to quantify the overall association between walking speed and search spread.

### Centroid angles and V-test

Centroid direction was quantified for each ant run based on the angular position of the search centroid relative to its starting point. For each run, the direction from the starting point to the centroid was converted to an angular measure (0° = North, clockwise) and plotted on a unit circle alongside the expected nest direction (221°). For each treatment, a one-sample circular V-test was performed to test whether centroid directions are non-uniform and specifically clustered around the expected nest direction.

### Angular deviation from expected fictive nest direction

Angular deviation from the expected nest direction was quantified for each ant run using the same centroids as described above. For each centroid, the absolute angular deviation (in degrees) from the expected nest direction (221°, north-clockwise reference frame) was calculated and summarized for each treatment. Differences between treatments, were assessed using a two-sample permutation test on the difference in mean angular deviation (10,000 permutations).

### Spatial distance to fictive nest

Spatial distance to the fictive nest was quantified for each ant run using the same centroids as described above. For each centroid, the Euclidean distance (in meters) to the fictive nest location (10 m at 221° in the same x–y reference frame) was calculated and summarized for each treatment. Differences between treatments were assessed using a two-sample permutation test on the difference in mean distance (10,000 permutations).

### Ethical note

As the experiments were carried out with ants, this work is not subject to any animal welfare restrictions. Nevertheless, we took care to prevent any possible suffering of the animals in all experiments.

## Author contributions

AB and MK designed study. AB, KB, and JK performed experiments. AB, MF, and KB analyzed data. AB wrote first draft of manuscript. All authors were involved in the final version of the manuscript.

## Data availability

All raw data are provided in the supplemental information.

